# Autonomic and respiratory components of orienting behaviors are mediated by descending pathways originating from the superior colliculus

**DOI:** 10.1101/2021.06.10.447470

**Authors:** Erin Lynch, Bowen Dempsey, Christine Saleeba, Eloise Monteiro, Anita Turner, Peter GR Burke, Andrew M Allen, Roger AL Dampney, Cara M Hildreth, Jennifer L Cornish, Ann K Goodchild, Simon McMullan

## Abstract

The ability to discriminate competing, external stimuli, and initiate contextually appropriate behaviors, is a key brain function. Neurons in the deep superior colliculus (dSC) integrate multisensory inputs and activate descending projections to premotor pathways responsible for orienting and attention, behaviors which involve adjustments to respiratory and cardiovascular parameters. However, the neural pathways that subserve the physiological components of orienting are poorly understood. We report that orienting responses to optogenetic dSC stimulation are accompanied by short-latency autonomic, respiratory and electroencephalographic effects in awake rats, closely mimicking those evoked by naturalistic alerting stimuli. Physiological responses were not accompanied by detectable aversion or fear and persisted under urethane anesthesia, indicating independence from emotional stress. Moreover, autonomic responses were replicated by selective stimulation of dSC inputs to a subregion in the ventromedial medulla containing spinally-projecting premotor neurons. This putative disynaptic pathway from the dSC represents a likely substrate for autonomic components of orienting.

## Introduction

The neural networks that control the circulatory and respiratory systems are co-opted by intrinsic and extrinsic stressors such as pain, fear and anger. Known widely as ‘defense’ or ‘fight-or-flight’ responses, this context-specific co-activation of cardiovascular, respiratory and behavioral states corresponds, in humans, with the conscious perception of emotion (Dampney et al., 2008; Kataoka et al., 2020; Silva et al., 2016). However, *defensive behaviors* are preceded in all vertebrates by *orienting responses*, the initial component of a pattern of activity that can transition to behaviors such as escape from a predator or pursuit of a prey, depending on the precise nature of the stimulus (Krauzlis et al., 2018).

In this context, novel, often innocuous, salient environmental stimuli that drive orienting also generate pronounced transient cardiovascular and respiratory effects which are an integral component of the classic orienting response. For example, unexpected sounds produce increases in arterial pressure (Baudrie et al., 1997; Caraffa-Braga et al., 1973; Rettig et al., 1986; Yu and Blessing, 1997), mesenteric and cutaneous vasoconstriction (Caraffa-Braga et al., 1973; Yu and Blessing, 1997), and increases in respiratory rate (Kabir et al., 2010; Nalivaiko et al., 2012), with little change in heart rate (Baudrie et al., 1997; Kabir et al., 2010; Nalivaiko et al., 2012). Visual or somatosensory stimuli can also evoke similar cardiovascular and respiratory responses (Kabir et al., 2010; Yu and Blessing, 1997). These effects are also typically accompanied by cortical electroencephalographic (EEG) desynchronization, superimposed by a prominent hippocampal theta rhythm (5-8 Hz) (Nalivaiko et al., 2012; Yu and Blessing, 1997), an electrophysiological signature of orientating, attention, and navigational behaviors, widely interpreted as an index of engagement with the environment (Buzsáki, 2006).

The superior colliculus (SC) is a phylogenetically conserved brain region that plays a critical role in generating innate behavioral responses, including motor components of orienting, to salient environmental stimuli (Isa et al., 2020; Sparks et al., 1990). Its deeper layers (dSC) receive convergent inputs from visual, auditory and somatosensory receptors, as well as from other brain regions including the basal ganglia and frontal and parietal cortices, and provide direct descending projections to motor and premotor nuclei in the brain stem and spinal cord that mediate behavioral responses (Boehnke and Munoz, 2008; Dean et al., 1989; May, 2006; Meredith and Stein, 1986). Under anesthesia, electrical or chemical stimulation of sites within the dSC also evokes physiological effects such as pupillary dilation, increases in arterial pressure, respiratory activity and sympathetic vasomotor nerve activity (Iigaya et al., 2012; Keay et al., 1990; Keay et al., 1988; Netser et al., 2010), whereas, dSC disinhibition augments sympathetic and respiratory responses to brief auditory, visual and somatosensory stimuli (Muller-Ribeiro et al., 2014; Muller-Ribeiro et al., 2016). Collectively, these observations suggest that, although normally subject to tonic inhibitory control, neurons in the dSC have the capacity to generate behavioral motor responses to alerting environmental cues and may also trigger autonomic and respiratory changes that support motor responses. However, the central pathways that underlie respiratory and autonomic components of dSC-mediated responses, and the behavioral contexts in which they are activated, are unclear.

Here we investigate the hypothesis that dSC neurons drive autonomic and respiratory adjustments independent of the pillars of the classic defense response: cortical processing, emotion and psychological stress. Using optogenetic stimulation in awake rats, we show that dSC stimulation evokes orienting behaviors that are accompanied by short-latency autonomic, respiratory and EEG effects that closely mimic responses evoked by naturalistic alerting stimuli. We show that these responses occur in the absence of detectable aversion or fear responses, indicating that they are not part of the defense response, and that such autonomic and respiratory effects are maintained under anesthesia and driven by a descending pathway that includes a relay in the ventromedial medulla oblongata.

## Results

### dSC stimulation drives alerting, orienting behavior, and physiological responses in awake rats

Experiments focused on a region of the rat caudolateral dSC (cl-dSC: Figure 1) which, when disinhibited, unmasks respiratory, cardiovascular and somatomotor responses to visual and auditory stimuli under urethane anesthesia (Muller-Ribeiro et al., 2014; Muller-Ribeiro et al., 2016). Following viral transduction of this region to express Channelrhodopsin2 (ChR2), optogenetic stimulation (63 – 160 mW/mm^2^, 5-10 Hz, 20 ms pulses, 5-10 s) evoked stimulus-dependent orienting behavior in awake rats, first detectable as a pause in ongoing activity with contraversive head-turning (i.e. turning in the direction contralateral to the side stimulated, Figure 1A). This response was reproducible within and between animals and was not seen in animals injected with a control, non-ChR2-expressing vector. As stimulus intensities increased, head-turning became more pronounced and sometimes led to transient full-body turning on the spot (Figure 1A, Video 1). At the highest stimulus intensities tested (320 mW/mm^2^, 10-20 Hz, 20 ms pulses, 5-10 s) the initial pause and head-turning component of the response was often bypassed, and instead rats transitioned into full-body turning on the spot, consistent with recent reports in the mouse and rat (Isa et al., 2020; Soper et al., 2016). Co-ordinated locomotor responses, such as running, were rare, and clear defensive behaviors, such as freezing, explosive escape or retreat, typical of stimulation of the rostromedial tectal defense pathway (Isa et al., 2020), were only seen in 1/33 rats.

**Figure 1.**
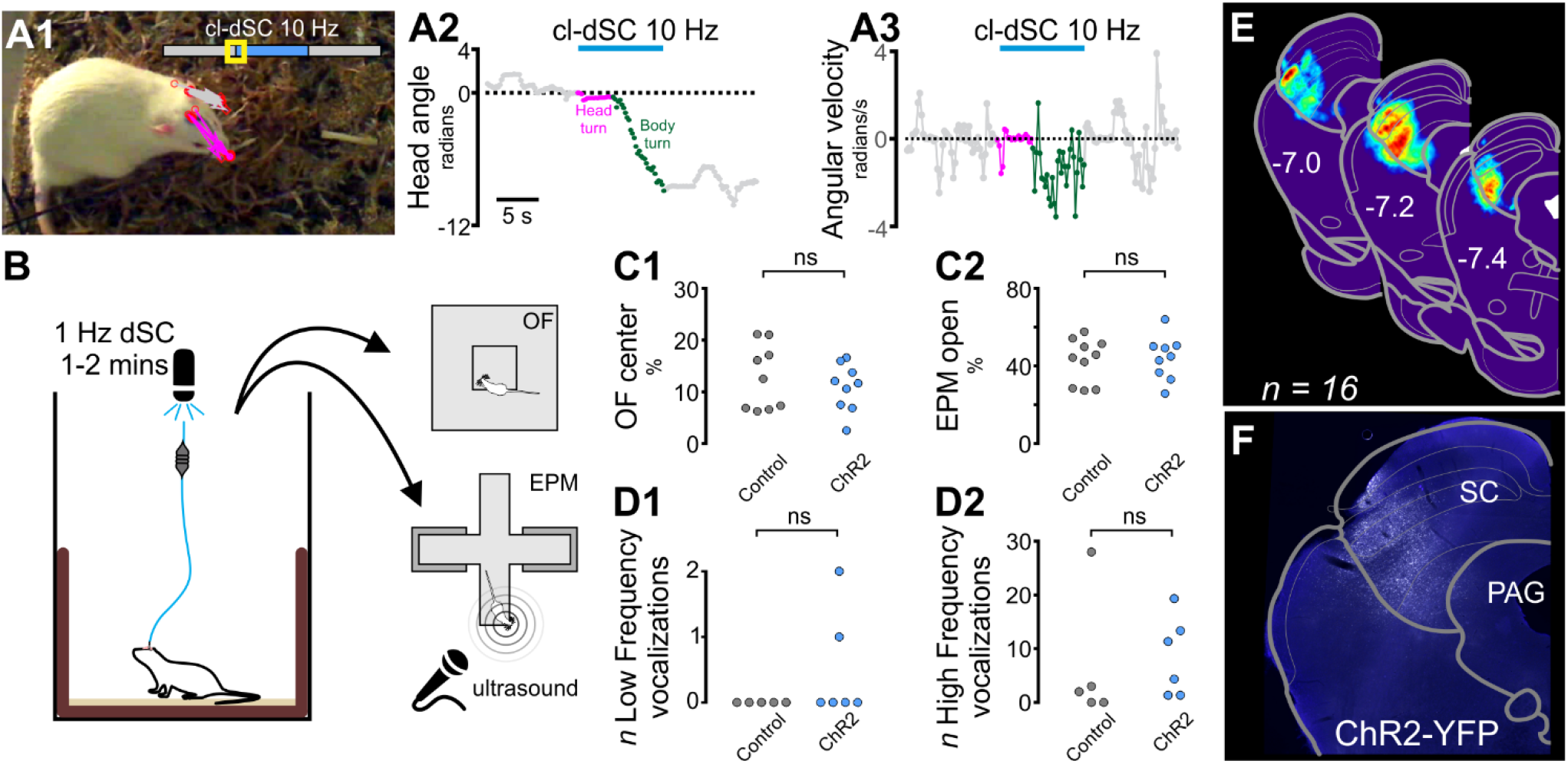
Caudolateral dSC stimulation evokes orienting responses but not anxiety-like behavior. A. Video-tracking shows typical response to moderate-intensity cl-dSC stimulation in ChR2-treated rat, characterized by immediate immobility and contraversive head-turning which transitions into full-body turning. A1 shows head position after 1 s of stimulation; head position in video frames denoted by the yellow box on the time scale is indicated by gray (baseline) and magenta (stimulation) arrows. A2-3 show head angle (with respect to orientation at the start of stimulation) and angular velocity during the same experiment. B. The presence of anxiety-like behaviors were assayed immediately after optical cl-dSC stimulation using the open field (OF, C1) or elevated plus maze (EPM, C2) apparatus: no significant difference was detected between rats treated with ChR2 or control vectors. Similarly no effects on incidence of low- or high-frequency ultrasonic vocalizations (USVs) were detected while exploring the EPM (D1 & 2). E. Distribution of normalized ChR2-YFP fluorescence in 16 animals; representative example shown in panel F. ns = not significant. Gray circles here and throughout indicate animals injected with control vectors; blue circles indicate animals injected with AAV-Syn-ChR2-YFP.

At the conclusion of cl-dSC activation, motor effects abruptly ceased and rats reverted to their previous behaviors without discernable signs of fear or heightened vigilance, such as aversion to approach or handling. To more closely assess whether stimulation of the cl-dSC elicited anxiety-like behaviors that outlasted stimulus presentation, we adapted the stimulus protocol (320 mW/mm^2^, 1 Hz, 20 ms pulses, 1-2 min: Figure 1B), after which rats were immediately transferred onto either an open field or an elevated plus maze apparatus. Under these conditions cl-dSC stimulation was not associated with detectable changes in open field (n = 9 ChR2 vs. 9 control rats, unpaired t-test, P = 0.6) or plus maze exploration (n = 10 ChR2 vs. 9 control rats, unpaired t-test, P >0.99, Figure 1C). Similarly, activation of the cl-dSC was not associated with low- or high-frequency ultrasonic vocalizations, indices of aversive and appetitive stimuli, respectively (Brudzynski, 2007; Brudzynski, 2013), during stimulation in the home cage (unpaired t-test, P = 0.2, data not shown, n = 6 ChR2 vs. 5 control rats) or exploration of the plus maze (unpaired P = 0.9, n = 6 ChR2 vs. 5 control rats, Figure 1D). On the other hand, 1 Hz cl-dSC stimulation continued to evoke orienting and motor effects, quantified as an increase in circling and distance travelled during stimulation, measured by overhead video tracking of the nose, without exerting measurable effects on cage exploration in the post-stimulation period (Figure 2). We conclude that optogenetic cl-dSC stimulation evokes acute, transient, stereotypic orienting-like motor responses, but that these are not associated with anxiety-like behaviors.

**Figure 2.**
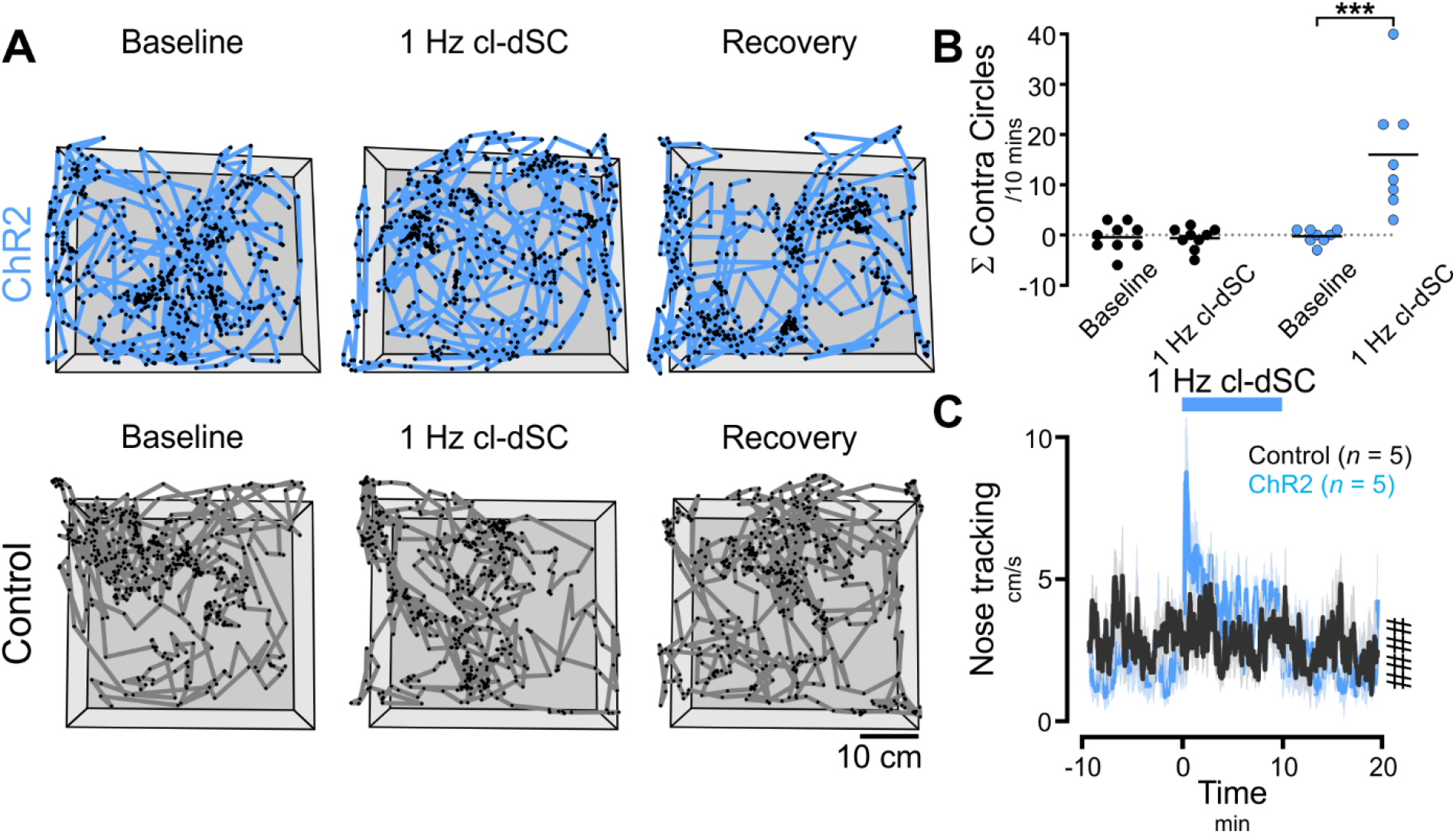
Behavioral responses to low frequency cl-dSC stimulation in rats injected with ChR2 and control vectors. A Examples of tracked nose position before, during and after 10 mins 1 Hz cl-dSC stimulation (position at 1s intervals denoted by black dots); in ChR2-treated animals cl-dSC stimulation was associated with contraversive circling (B) and increased distance travelled (C), particularly during the first two minutes of stimulation. Gray boxes in A denote floor and walls of home cage; ***: P<0.001 paired t-test, ####: interaction P<0.0001, 2-way RM ANOVA. SEM here and elsewhere indicated by lighter shading.

Much like behavioral responses to naturalistic salient stimuli (Nalivaiko et al., 2012), orienting responses evoked by cl-dSC stimulation were accompanied by profound transient elevations in respiratory rate and EEG desynchronization together with increased theta power (5-8 Hz), even at stimulation intensities that evoked minimal motor effects (63 – 160 mW/mm^2^, 5-10 Hz, 20 ms pulses, 10 s, head turning only, Figure 3A). In the rat, orienting responses to innocuous naturalistic stimuli are associated with transient tail vasoconstriction (de Menezes et al., 2009; Nalivaiko et al., 2012), resulting in tail cooling and core heat retention. Infrared tail thermography revealed similar effects in response to cl-dSC stimulation (320 mW/mm^2^, 1 Hz, 20 ms pulses, 10 min): tail temperature dropped to a nadir of −2.0 ± 0.4 °C during photostimulation and then rebounded to a peak of +1.1 ± 0.4°C relative to baseline over the ten minutes after stimulation (RM 2-way ANOVA interaction P <0.0001, Figure 3B). Activation of respiratory and autonomic outputs is therefore an inherent component of orienting responses evoked by cl-dSC stimulation.

**Figure 3.**
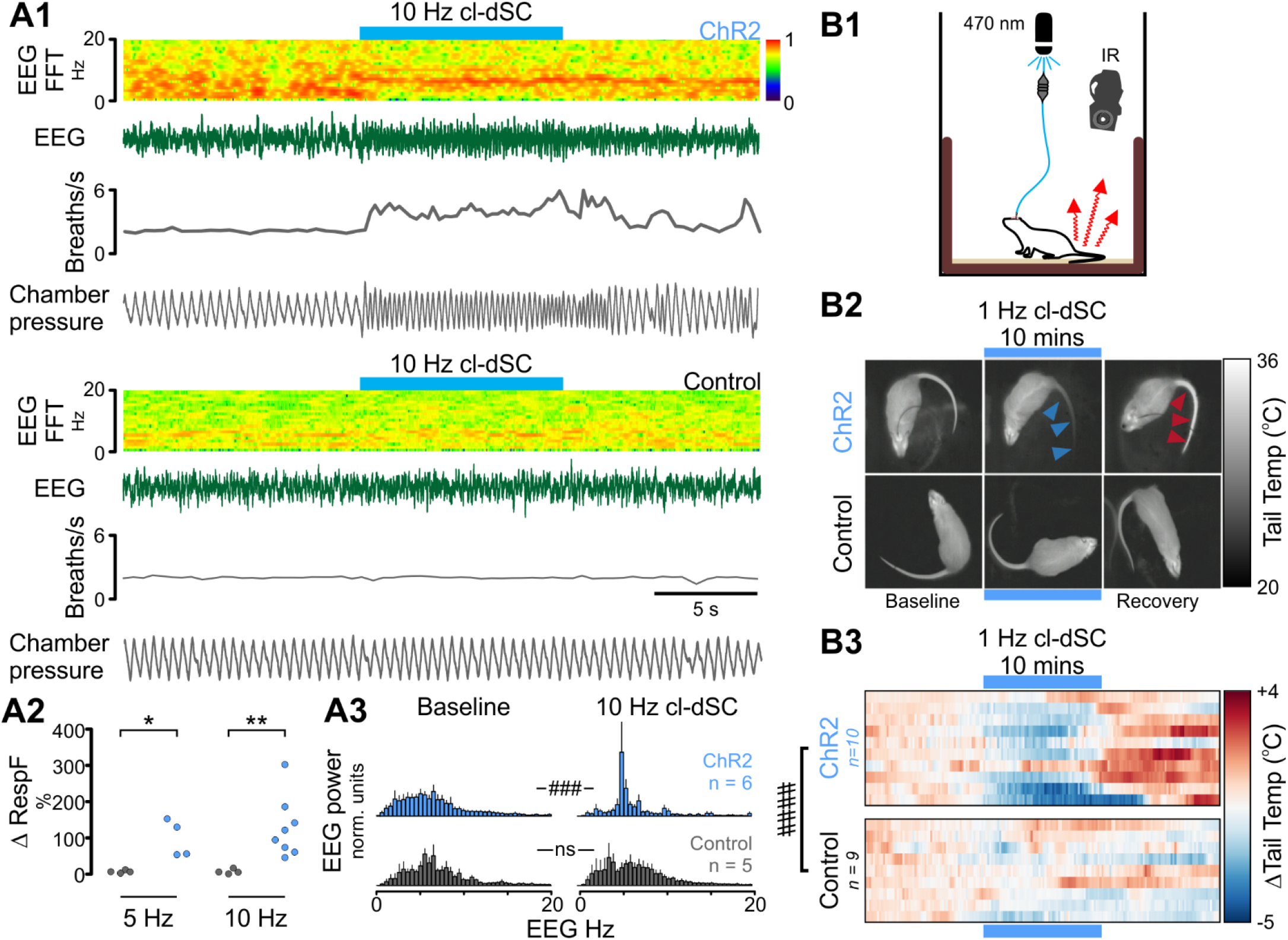
cl-dSC stimulation drives arousal, ventilation, and tail vasoconstriction in ChR2-treated but not in vector control rats. A1. Simultaneous recording of electroencephalogram (EEG) and respiratory rate from ChR2 (upper) and control vector treated (lower panels) animals showing tachypnoea and EEG desynchronization (0.5 - 4.5 Hz) with increased theta power (5-8 Hz). Pooled data shown in A2 (respiratory rate) and A3 (EEG frequency). B1. Tail temperature was measured by infra-red thermography before, during and after 10 minutes of 1 Hz cl-dSC stimulation. B2. Thermal imagery illustrates tail-cooling in ChR2 (blue arrowheads) experiment during stimulation, and tail temperature rebound (red arrowheads) during the recovery period. B3. Heat maps illustrate recordings from 10 ChR2 and 8 control vector treated rats: each row represents an individual experiment. *, **: Mann Whitney P<0.05, 0.01, ###, ####: 2-way RM ANOVA interaction P<0.001, <0.0001.

### Physiological responses to cl-dSC stimulation are maintained under anesthesia

Adjustments to blood pressure, sympathetic nerve activity and breathing are integral features of motor behaviors that result from both peripheral sensory signals and feedforward from hypothalamic central command circuits (reviewed by Dampney et al., 2008). To differentiate physiological responses evoked by cl-dSC stimulation from those dependent on motor components of orienting responses, recordings of respiratory, blood pressure, heart rate and postganglionic splanchnic sympathetic nerve responses to cl-dSC stimulation were made in urethane-anesthetized rats (Figure 4). Low-frequency stimulation (320 mW/mm^2^, 0.5 Hz, 20 ms pulses, 200–600 repeats) revealed constant-latency sympathoexcitatory responses in 8/11 cases (onset 43 ± 3 ms, peak 101 ± 3 ms, Figure 4D). In most experiments where sympathetic responses were observed, intensity-dependent respiratory and cardiovascular responses to short trains of repetitive cl-dSC stimulation were also observed (320 mW/mm^2^, 10-20 Hz, 20 ms pulses, 10 s). Responses were characterized by abrupt increases in respiratory frequency and respiratory burst size (Figure 4C, 8/8 cases), increased mean splanchnic sympathetic nerve activity (SNA, Figure 4E, 6/8 cases), and systolic arterial blood pressure (sAP, Figure 4F, 7/8 cases), with variable effects on heart rate (Figure 4G).

**Figure 4.**
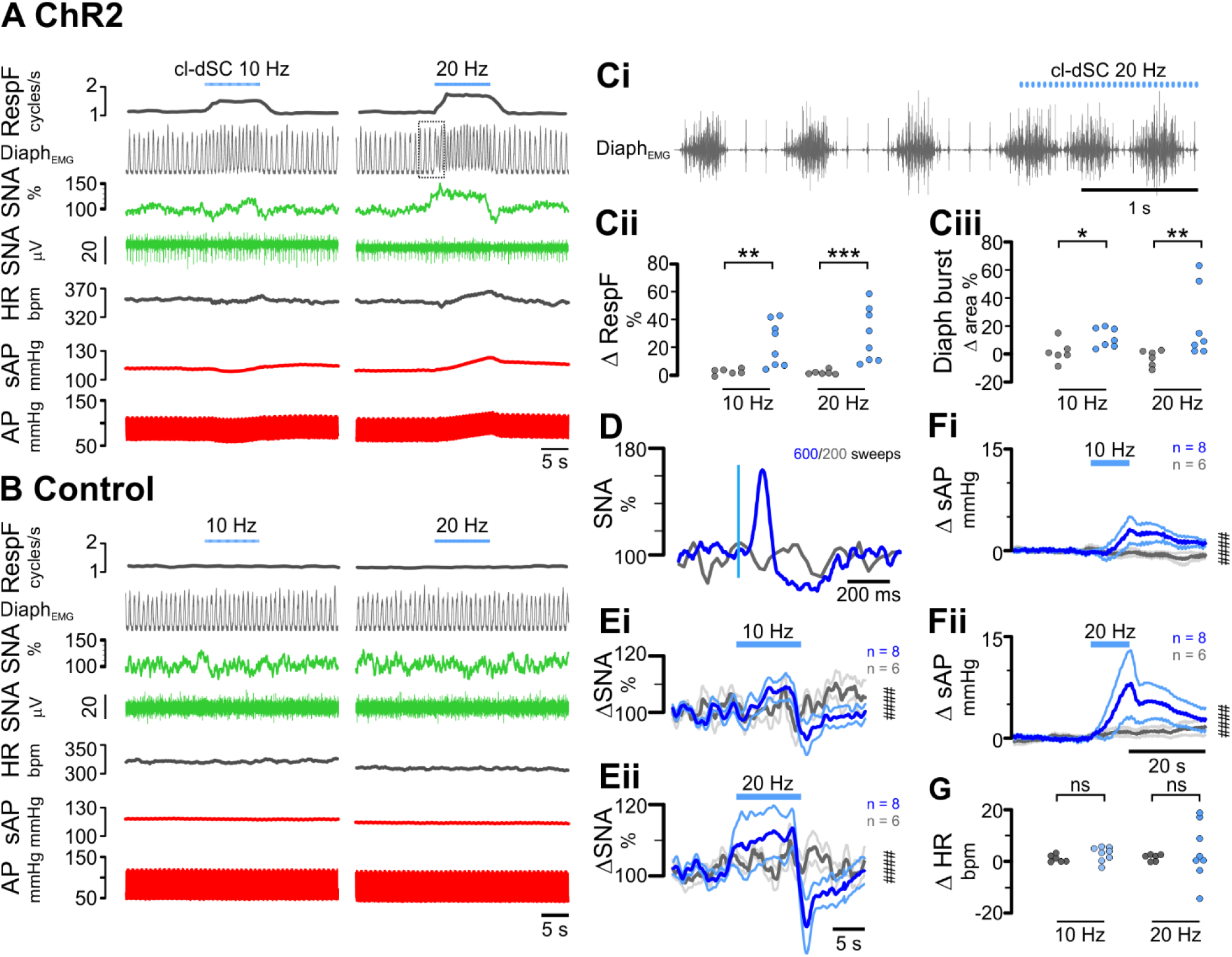
Respiratory and cardiovascular effects of cl-dSC stimulation under urethane anesthesia. Representative recordings from rats treated with ChR2 (A) and control (B) vectors. Boxed region in Panel A denotes expanded trace shown in Panel Ci. Respiratory effects (Panel C) manifested as an increase in diaphragmatic EMG burst frequency (Cii) and amplitude (Ciii). Single-pulse cl-dSC stimulation evoked short-latency monophasic bursts in SNA (D: vertical blue line denotes laser onset); repetitive cl-dSC stimulation evoked frequency-dependent recruitment of SNA (E) and systolic blood pressure (sAP, F), with variable effects on heart rate (HR: G). Bold traces in E & F indicate mean, light traces indicate SEM. *, **, ***: Mann Whitney P<0.05, <0.01, <0.001; ####: 2-way RM ANOVA interaction P<0.0001.

### Physiological responses to cl-dSC stimulation are mediated via relay in the medullary reticular formation

The short latency of splanchnic sympathetic responses to cl-dSC stimulation, and our previous observation that cl-dSC-facilitated autonomic responses to sensory stimulation persist following pre-collicular decerebration (Muller-Ribeiro et al., 2014), suggest a direct descending neural pathway from the cl-dSC that does not involve ascending loops through forebrain limbic or hypothalamic defense regions. Using anterograde tract tracing, we identified putative connections between cl-dSC output neurons and hindbrain autonomic and respiratory relays. Microinjection of AAV-CBA-tdTomato into the cl-dSC revealed a clearly defined ascending projection to the thalamus and hypothalamus, a projection to the contralateral superior and inferior colliculi, via the commissural nucleus of the inferior colliculus, and a descending projection to the brainstem (Figure 5S1). Confocal imaging of tdTomato-labelled hindbrain terminal fields identified close appositions in candidate autonomic nuclei (Figure 5S2), including moderate innervation of catecholaminergic neurons in the A5, A6 (locus coeruleus) and A7 (parabrachial/Kölliker-Fuse) regions. A more conspicuous and largely ipsilateral nexus of fibers and arborized varicosities extended across subnuclei within the ventromedial medulla to encompass the gigantocellular alpha (GiA) and paragigantocellular reticular nuclei and obscurus, pallidus, and magnus raphé nuclei (Figure 5A & B).

**Figure 5.**
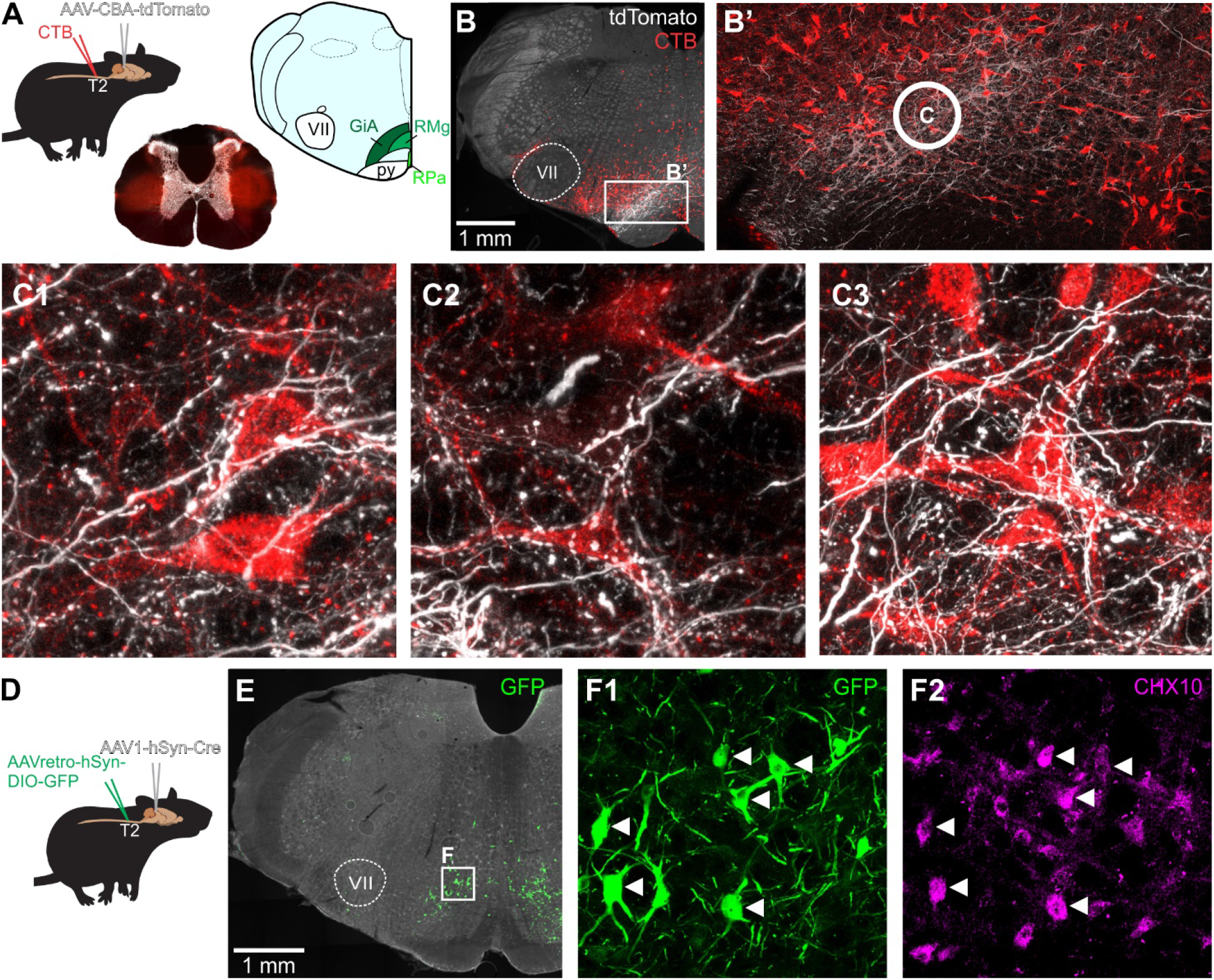
cl-dSC projections to spinally projecting neurons in the ventromedial medulla: A. Tracing strategy: AAV-CBA-tdTomato was used for anterograde labelling of cl-dSC neurons (white channel in B & C); microinjection of CTB at the thoracic intermediolateral cell column was used for retrograde labelling of bulbospinal neurons (red channel, brightfield photomicrograph in A shows CTB injection sites). cl-dSC injections resulted in extensive and predominantly ipsilateral terminal field labelling in the ventromedial medulla (B), dorsal to the pyramidal tract (py), spanning the region from the midline raphe magnus (RMg) to the lateral gigantocellular reticular nucleus (GiA: B’). C. High-power confocal images of CTB-labelled GiA neurons from the circled region in B’ in close apposition to cl-dSC terminals. D. Similar results were obtained using Cre-mediated trans-synaptic tagging: AAV1-hSyn-Cre injection in the cl-dSC results in transport of Cre to post-synaptic targets. Spinally projecting neurons were transduced by injection of AAVretro-hSyn-DIO-GFP in the thoracic spinal cord. GFP expression is thus limited to spinally projecting neurons that receive inputs from the cl-dSC: GFP-immunoreactive GiA neurons are shown in E. F. Confocal images from the boxed region in E show colocalization of GFP with CHX10 immunoreactivity (arrowheads).

**Figure 5 S1.**
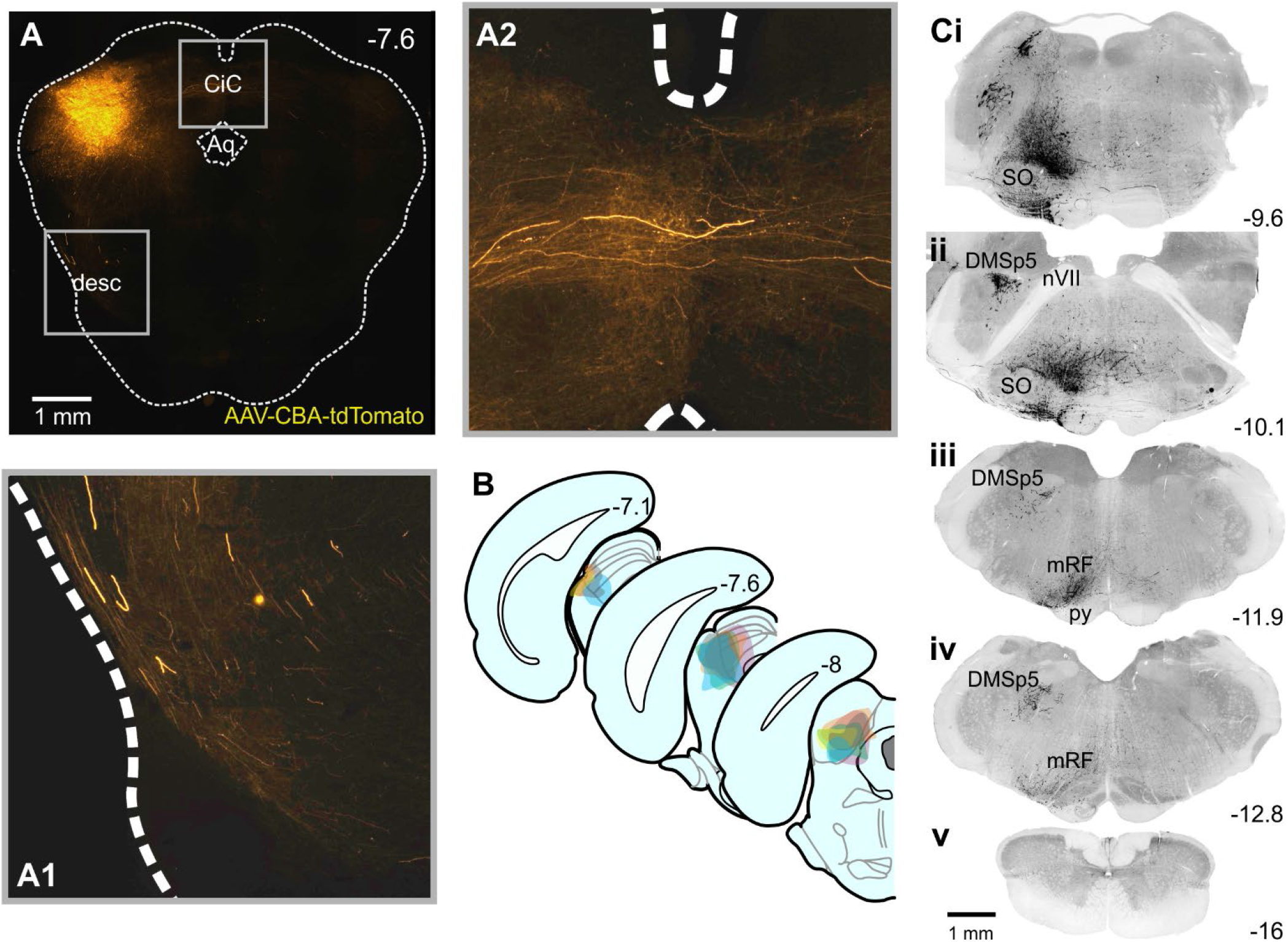
Anterograde labelling from the cl-dSC: A. Example of AAV-CBA-tdTomato injection site in the cl-dSC. B. Composite diagram showing distribution of injection sites in 6 experiments. C. Shows histological sections from one experiment, illustrating the brainstem distribution of tdTomato labelling. Fibers emerging from the injection site followed either a descending trajectory via the lateral midbrain (desc, A1), or to the contralateral colliculi via the commissure of the inferior colliculus (CiC, A2). The descending bundle split into two tracts in the pons, a more substantial ventral bundle which emerged in the brainstem, surrounding the superior olive on all sides (SO, Ci), and a lesser and exclusively ipsilateral dorsal tract that innervated the dorsomedial spinal trigeminal nucleus (DMSp5, Cii) with large caliber fibers. The ventral branch occupied a column extending from the ventral surface to an apex at locus coeruleus (see Figure 5S2, panel C), was sparsely mirrored on the contralateral brainstem, and comprised both large and fine caliber fibers and putative terminals. Caudal to the facial nerve (nVII, Cii) the ventral branch was most densely concentrated in the medullary reticular formation (mRF, Ciii) dorsal to the pyramidal tract (py) at the level of the caudal pole of the facial nucleus. Labelled fibers were infrequently encountered in the caudal medulla (Civ) and virtually absent in spinal sections (Cv). Coordinates indicate rostrocaudal position with respect to Bregma.

**Figure 5S2:**
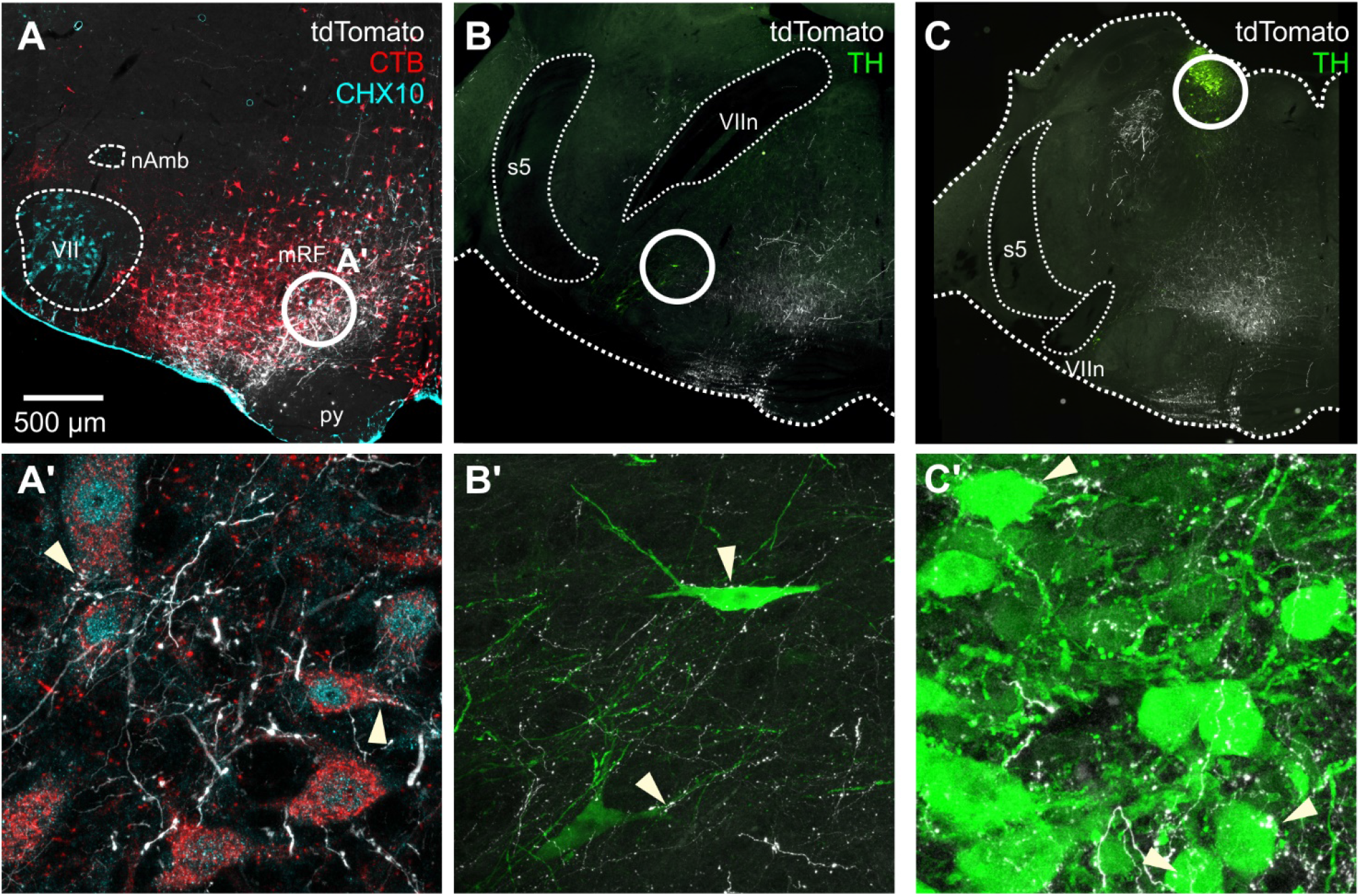
Phenotypes of putative targets of cl-dSC outputs, labelled by AAV-CBA-tdTomato: top row identifies region of interest (circled), lower panels highlight putative close appositions with CHX10 bulbospinal neurons in the mRF (identified by retrograde transport of CTB: A) and tyrosine hydroxylase (TH) immunoreactive neurons in the A5 (B), and A6 (C) cell groups.

Spinally projecting neurons in this region, including V2a neurons that express the transcription factor CHX10, are considered key players in the initiation and arrest of locomotion and orienting behaviors (Dougherty and Kiehn, 2010; Usseglio et al., 2020), and were confirmed recipients of putative inputs from labelled cl-dSC neurons (Figures 5C & F, 5S2), with the caveat that not all close appositions identified under light microscopy are functional synapses (reviewed by Saleeba et al., 2019). We therefore used an intersectional viral tracing strategy, based on the anterograde trans-synaptic trafficking of Cre recombinase when expressed by the AAV1 serotype (Zingg et al., 2017), to bypass these limitations and selectively label spinally projecting neurons that receive synaptic input from the cl-dSC. In these experiments AAV1-hSyn-Cre was injected in the cl-dSC to control GFP transcription in post-synaptic spinally projecting neurons, transduced by injection of the Cre-dependent retrograde vector AAVretro-hSyn-DIO-GFP at the intermediolateral column of the T2 spinal cord (Figure 5E). This approach recapitulated the findings of our conventional anterograde tracing experiments, revealing GFP-immunoreactive GiA neurons in 3 of 4 rats (Figure 5F) of which also 38% were found to be CHX10 immunoreactive (14/38 neurons, 4 confocal stacks from one animal, Figure 5G). No GFP+ neurons were identified in any other region except for a small number of neurons in the sub-coeruleus region of the pontine reticular nucleus (n=1/4 rats). These findings provide evidence for a disynaptic pathway that links the cl-dSC to thoracic spinal outputs via a relay in the GiA.

Populations of neurons in this region of the rostral ventromedial medulla form monosynaptic connections with spinal motoneurons (Esposito et al., 2014) and have recently been implicated in coordinating motor effects of dSC stimulation (Isa et al., 2020). However, the GiA is functionally diverse; it is a major site of sympathetic premotor neurons that innervate vasomotor and non-vasomotor targets (reviewed by Dampney, 1994; Stornetta, 2009), and its neurochemically diverse constituents are also implicated in respiratory and somatosensory functions (Crone et al., 2012; Depuy et al., 2011; Morris et al., 1996; Porreca et al., 2002), suggesting a plausible role for this pathway in mediating physiological responses to cl-dSC stimulation. To investigate this possibility, we injected AAV-Syn-ChR2-YFP in the cl-dSC and optically stimulated ChR2-transduced axon terminals in the GiA in urethane-anesthetized rats. Autonomic and respiratory responses were consistent with those observed following cl-dSC stimulation, and were only observed in rats in which expression of ChR2-YFP in the ipsilateral GiA was histologically verified (Figure 6). The profiles of splanchnic sympathetic (Figure 6C) and pressor (Figure 6D) responses to 20 Hz GiA stimulation were not significantly different from those evoked by 20 Hz cl-dSC stimulation (2-way ANOVA interaction P>0.99 for both sympathetic and pressor responses, n = 7, direct comparison in Figure 6S1A). On the other hand, although GiA stimulation increased respiratory frequency in 4/7 cases, effects were consistently smaller than those evoked by cl-dSC stimulation (10 ± 3 vs. 29 ± 7%, unpaired t-test P = 0.036, Figure 6S1B). In 4/5 animals tested, low-frequency GiA stimulation (320 mW/mm^2^, 0.5 Hz, 20 ms pulses) evoked sympathoexcitatory responses that occurred at a shorter latency (onset 30 ± 1 vs. 43 ± 3 ms, unpaired t-test P = 0.02) but equivalent peak amplitude (151 ± 14 vs. 163 ± 25%, unpaired t-test P = 0.76) compared to cl-dSC stimulation (Figure 6E).

**Figure 6.**
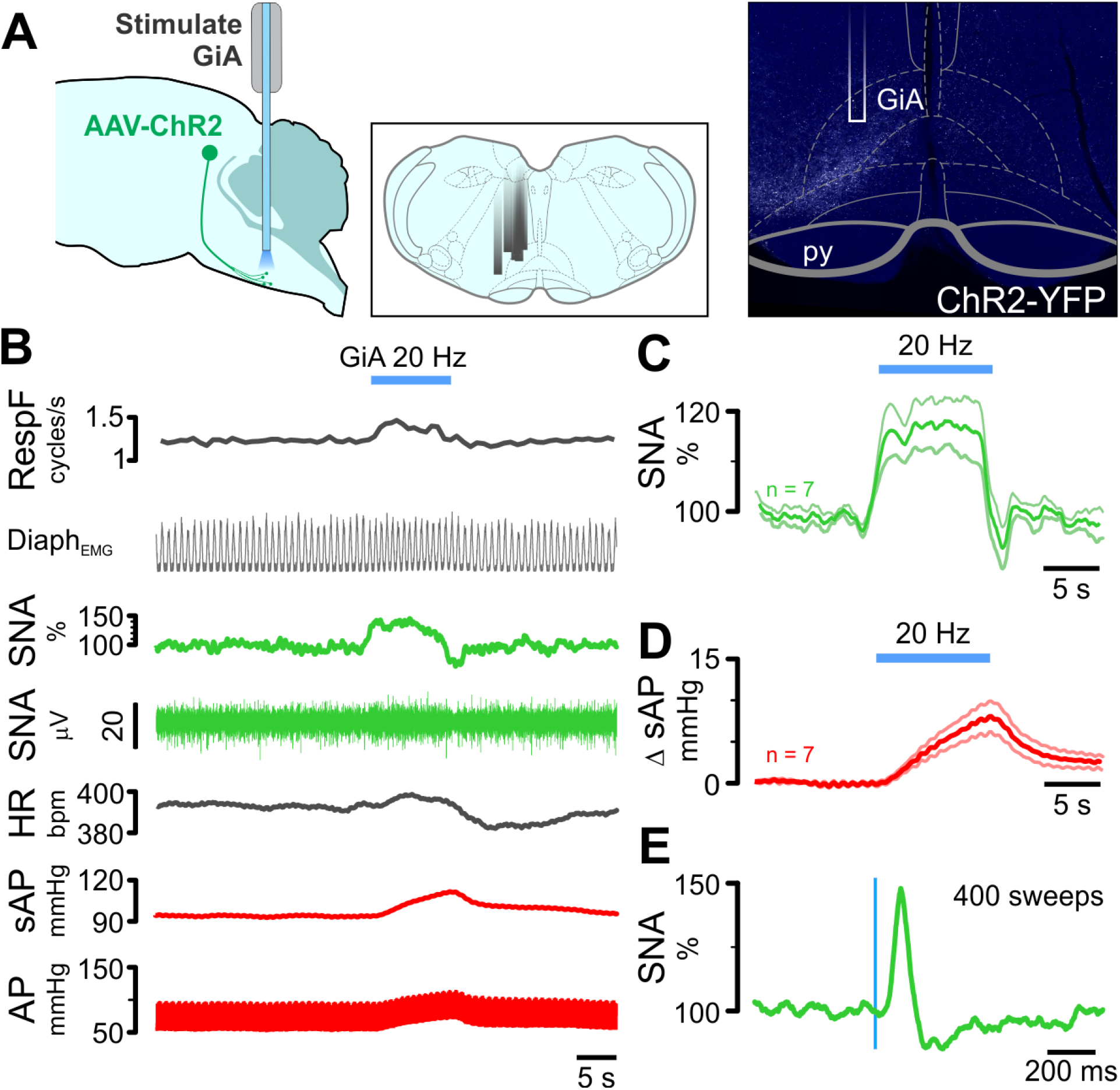
Respiratory and cardiovascular responses to optogenetic stimulation of cl-dSC-GiA terminals under urethane anesthesia. Panel A illustrates (L-R) experimental strategy, locations of fiber optic cannulae in different experiments, and example illustrating cannula position and terminal ChR2 labelling. B. Physiological recording of respiratory, sympathetic and cardiovascular responses to GiA stimulation; pooled sympathetic and pressor responses from 7 experiments are shown in C & D. E. Shows stimulus-triggered average of sympathetic response to 0.5 Hz stimulation from the experiment shown in B.

**Figure 6 S1.**
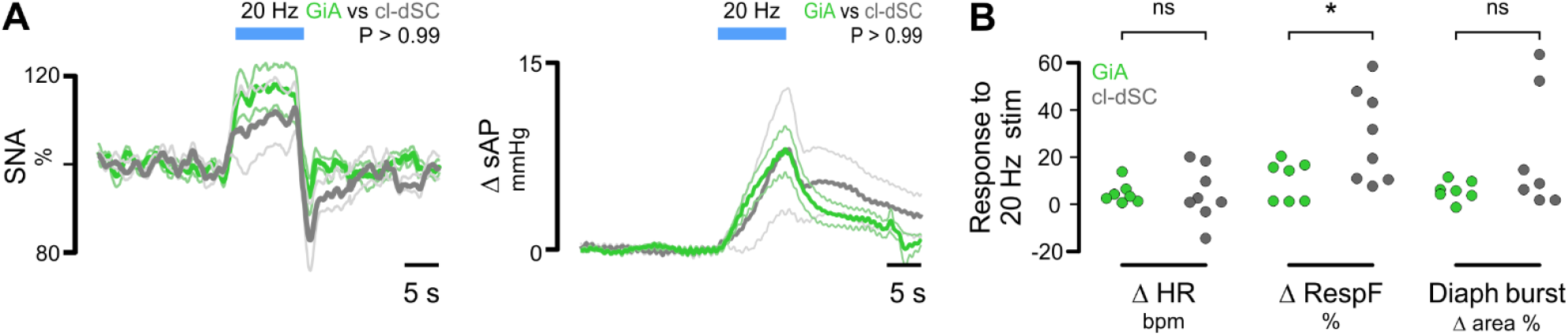
Direct comparison of responses evoked by optogenetic stimulation of cl-dSC neurons (dark gray) and their GiA terminals (green). Outputs examined were sympathetic nerve activity & systolic blood pressure (A), and heart rate, respiratory frequency and respiratory burst area (B). Significantly different responses were only detected in respiratory frequency (*: P<0.05, unpaired t-test).

Extracellular recordings confirmed that spanchnic sympathetic responses to GiA stimulation are likely mediated via direct activation of local sympathetic premotor neurons (Figure 7). In two urethane-anesthetized rats, 7/7 spontaneously active (12.6 ± 5.1 Hz) spinally-projecting GiA neurons (spinal conduction velocity 12.5 ± 2.8 m/s) exhibited frequency-dependent excitatory responses to cl-dSC stimulation (320 mW/mm^2^, 20 ms pulses: 10 Hz: +28 ± 9%; 20 Hz: +72 ± 14%, Figure 7C), and low frequency cl-dSC stimulation (320 mW/mm^2^, 0.5 Hz, 20 ms pulses) revealed short latency excitatory responses in 2/4 cells tested (8, 17 ms Figure 7D). Several units (4/7) were identified as putative sympathetic premotor neurons based on entrainment of unit discharge to blood pressure pulsatility (n = 3, Fig 7E) or covariation of neuronal discharge with SNA, revealed by spike-triggered averaging (n = 4, data not shown).

**Figure 7.**
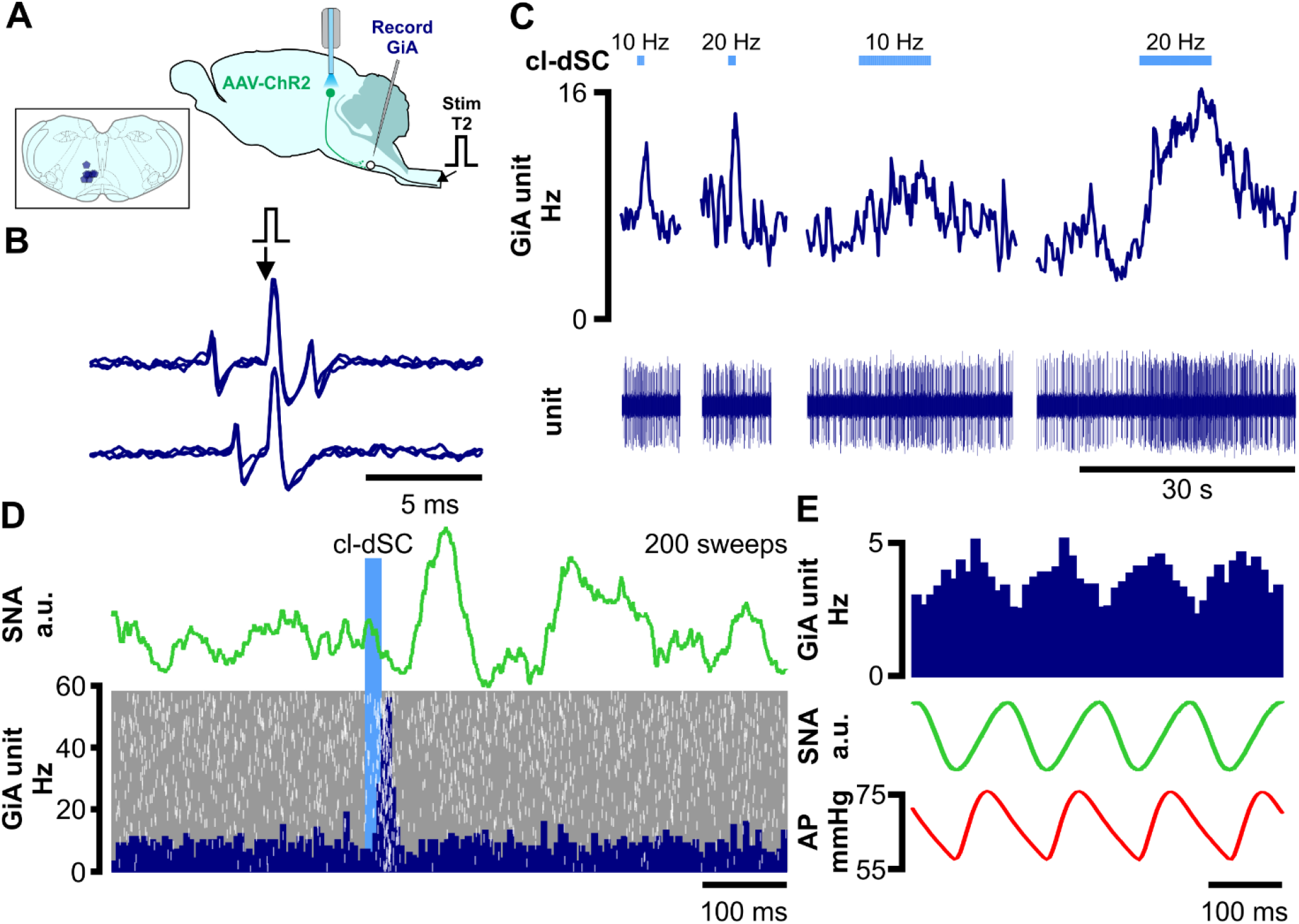
Optogenetic cl-dSC stimulation activates spinally projecting cardiovascular and non-cardiovascular GiA neurons. A. Extracellular recordings were made from bulbospinal GiA neurons under urethane anesthesia after transduction of cl-dSC neurons with ChR2 and fiber optic implantation; inset shows recording sites of 7 neurons. B. All spinally projecting neurons exhibited constant-latency antidromic spikes in response to electrical stimulation of the T2 spinal cord (top trace) which collided with spontaneous orthodromic spikes (lower trace). C. cl-dSC stimulation evoked frequency-dependent increases in baseline activity: trace shows typical responses to short (1s) and long (10 s) trains of stimulation at 10 and 20 Hz (same neuron as B). D. Low-frequency cl-dSC stimulation evoked short-latency response in the same neuron (laser-triggered peristimulus time histogram with overlaid raster, lower panel) that preceded simultaneously recorded sympathoexcitation (green trace, upper panel). E. Putative sympathetic premotor neurons were identified by covariation of spontaneous neuronal activity (top trace) with SNA (green) and pulsatile arterial pressure (lower trace).

## Discussion

Here we show that, in freely behaving rats, stimulation of the cl-dSC at intensities that drive orienting responses, EEG desynchronization and theta rhythm, evokes transient respiratory and cutaneous vasomotor responses (tail temp) that are consistent with the effects evoked by natural salient stimuli (de Menezes et al., 2009; Nalivaiko et al., 2012; Yu and Blessing, 1997). The brief nature of responses and the absence of anxiety-like or avoidant behaviors differentiates them from autonomic and respiratory responses evoked by psychological stress, which outlast stimulus presentation (Ootsuka et al., 2017; Vianna et al., 2008; Vianna and Carrive, 2005), and suggests independence from the canonical defense response.

Detailed physiological recordings under anesthesia recapitulated the respiratory effects seen in behaving animals and revealed excitatory effects on splanchnic sympathetic nerve activity which were similar in latency to those evoked by direct stimulation of sympathetic premotor neurons (Abbott et al., 2009; Menuet et al., 2014). Stimulation of the cl-dSC also evoked pressor effects with little change in heart rate, mimicking the pattern of cardiovascular effects recruited by alerting stimuli, thought to result from activation of vasomotor sympathetic nerves and co-activation of cardiac sympathetic and parasympathetic nerves (Casto and Printz, 1990; Nalivaiko et al., 2012). Viral anterograde tracing of cl-dSC neurons defined two main descending pathways, a major ventral pathway to the ventromedial medulla, with evidence of monosynaptic connections with CHX10 and non-CHX10 bulbospinal neurons, and a minor dorsal pathway that targeted the dorsomedial spinal trigeminal nucleus. Stimulation of cl-dSC efferent terminals in the GiA region of the ventromedial medulla mimicked the autonomic pattern evoked by direct stimulation of the cl-dSC. Interestingly, the respiratory response to stimulation of dSC-GiA axons was reduced compared to direct stimulation of the cl-dSC, indicating that involvement of a separate supplementary respiratory circuit might be required for full elaboration of respiratory effects. Spinally projecting autonomic neurons in the same area were functionally identified *in vivo* by their transmission of cardiovascular information (Chen and Toney, 2010; McMullan et al., 2007; Morrison et al., 1988; Turner et al., 2013), and shown to receive excitatory input from the cl-dSC.

Previous studies reported that autonomic responses to visual, acoustic and somatosensory stimuli, unmasked by disinhibition of the cl-dSC, persist following pre-collicular decerebration (Muller-Ribeiro et al., 2014). We reasoned that the physiological effects of dSC stimulation may reflect a previously unrecognized descending pathway, and used anterograde tracing to identify putative relays to medullary respiratory and autonomic nuclei. We observed no anatomical evidence of innervation of the rostral ventrolateral medulla (RVLM) by cl-dSC output neurons. This is perhaps surprising, as RVLM neurons are: a major source of excitatory drive to sympathetic nerves and play a critical role in subserving a wide range of cardiovascular reflexes (Guyenet, 2006); implicated in mediating sympathetic responses to a range of physiological and psychological stressors (Abe et al., 2017; Guyenet et al., 2013; Zhao et al., 2017); and reported to receive monosynaptic input from the SC (Dempsey et al., 2017; Stornetta et al., 2015). In contrast, we found extensive arborization of cl-dSC efferents across multiple medullary reticular formation (mRF) compartments, most prominently the GiA, which is a major hub for descending control to spinal locomotor circuits (Bouvier et al., 2015; Dougherty and Kiehn, 2010; Kim et al., 2017; Perreault and Giorgi, 2019; Usseglio et al., 2020). Similar SC innervation of the mRF has been described previously in the mouse and rat (Isa et al., 2020; Redgrave et al., 1987; Usseglio et al., 2020). GiA neurons are potently stimulated by alerting stimuli (Furlong et al., 2014) and are implicated in mediating motor responses to SC stimulation (Isa and Sasaki, 2002). However, in addition to its motor functions, the GiA and lateral paragigantocellular nucleus contain abundant sympathetic premotor neurons that innervate brown adipose, visceral and cutaneous targets (Cano et al., 2001; Card et al., 2011; Morrison, 1999; Rathner and McAllen, 1999; Standish et al., 1995; Strack et al., 1989; Varner et al., 1992), making this region uniquely suited for integration of motor and autonomic commands from the SC. Notably, Cre-mediated trans-synaptic tagging of spinally projecting neurons failed to identify any other plausible direct relays between the cl-dSC and spinal sympathetic outputs as an alternative pathway for the short latency sympathetic responses observed.

Optogenetic stimulation of cl-dSC-GiA terminals evoked pressor and sympathetic effects equivalent to those elicited by cl-dSC activation, with sympathoexcitatory responses to single-pulse GiA stimulation occurring at shorter latencies than those evoked by cl-dSC stimulation, consistent with a role for this pathway in direct activation of medullary sympathetic premotor neurons. Extracellular recordings from spinally projecting GiA neurons, including those whose spontaneous activity correlated with sympathetic nerve activity, confirmed excitatory input from cl-dSC stimulation at a latency (~10 ms) that aligns well with the difference between the latencies of sympathoexcitatory responses evoked by cl-dSC vs. GiA stimulation (~13 ms). Taken together, these findings suggest that sympathetic responses to cl-dSC stimulation are principally due to activation of GiA neurons and confirm a role for bulbospinal sympathetic premotor GiA neurons in this pathway. Based on the latency of responses and the results of our anterograde tracing experiments, this pathway is likely disynaptic and unlikely to involve supramedullary sites.

In contrast, respiratory responses to cl-dSC-GiA terminal stimulation were consistently smaller than those evoked by direct cl-dSC stimulation, suggesting the necessary involvement of neurons within and beyond the GiA for full respiratory effects. Potential local mediators of respiratory responses include serotonergic raphe obscurus neurons and non-serotonergic GiA neurons, stimulation of which evokes tachypnoea (Depuy et al., 2011; Verner et al., 2004). Supramedullary contributors to cl-dSC-evoked tachypnoea could include the parabrachial/Kölliker-Fuse complex and periaqueductal grey, stimulation of which increases respiratory rate under anesthesia (Chamberlin and Saper, 1994; Hayward et al., 2003) and both of which were identified as sites of dSC terminal fields in the current study.

The present study identifies a novel pathway that links neurons in the SC with autonomic and respiratory actuators in the medulla, but two key questions remain. First, do cl-dSC projections target separate populations of GiA neurons that drive distinct descending motor and autonomic pathways, or do they target multifunctional GiA neurons that provide divergent excitatory drive to respiratory, sympathetic and somatic motor pools? The latter seems possible: genetic deletion of V2a neurons, the quintessential reticulospinal premotor cell group and a major dSC target (Dougherty and Kiehn, 2010; Usseglio et al., 2020), is associated with motor deficits (Crone et al., 2008; Crone et al., 2009) and a severe hypoventilation phenotype that persists in the isolated medullary slice (and therefore not attributable to spinal V2a depletion: Crone et al., 2012). These functional data are strongly supported by anatomical evidence: polysynaptic viral tracing indicates that at least 20% of GiA bulbospinal neurons provide dual innervation to motor and autonomic targets (Kerman, 2008). These findings are consistent with an emerging view of mRF neurons as containing distinct functionally diverse subpopulations (Usseglio et al., 2020), and would make them well-placed to serve as command modules for orchestration of adaptive motor, respiratory and autonomic responses to salient environmental cues.

The second question regards the behavioral contexts under which this pathway is activated. The superior colliculus is a critical node for the selection and execution of innate behavioral responses to multimodal environmental cues (Saidel, 2009; Yilmaz and Meister, 2013). Its structure is highly conserved across vertebrate groups (Butler and Hodos, 2005), and organized to reflect the topography of the sensory inputs that converge upon it (Stein and Meredith, 1993). As a consequence, different regions of the SC process inputs that vary not only in their somatotopy, but also in their ecological saliency (Comoli et al., 2012), and correspondingly evoke distinct patterns of motor activity, including orienting, predation, avoidance and escape (Basso and May, 2017; Furigo et al., 2010; Shang et al., 2019; Sparks et al., 1990; Yilmaz and Meister, 2013). Our experiments confirm a recent report by Isa et al. (2020), which describes intensity-encoded orienting-like behavior in response to optogenetic activation of the analogous region of the mouse caudolateral dSC, and contrast with the escape and freezing behaviors evoked by stimulation of the rostromedial dSC (Sahibzada et al., 1986; Wei et al., 2015). Whether the cl-dSC pre-autonomic pathway described here is implicated in more complex behaviors, or whether analogous descending pathways emerge from other regions of the dSC (perhaps supporting physiological components of other SC-mediated innate behaviors) remains to be seen. Like the cl-dSC, the rostromedial SC contains visually responsive neurons that project to the ipsilateral mRF, many of which are also activated by auditory and somatosensory stimuli (Isa et al., 2020; Westby et al., 1990). However, it is not yet clear whether descending efferents from the medial dSC contribute to respiratory or autonomic pathways: whereas early studies found equivalent pressor responses to electrical stimulation of the medial and lateral dSC (Keay et al., 1988), subsequent pharmacological studies found reactive sites were concentrated in the lateral dSC (Iigaya et al., 2012; Keay et al., 1990; Muller-Ribeiro et al., 2014).

In summary the results of the current study demonstrate that the caudolateral dSC co-activates motor and physiological outputs that mimic those observed during alerting and orienting behaviors. These responses are largely reproduced by stimulation of dSC axons in the mRF, a critical relay nucleus for driving motor responses in SC-mediated orienting. In driving autonomic components of descending orienting pathways, this disynaptic pathway potentially primes physiological systems for future phases of arousal/flight or fight behavior. The results of the present study, together with those Muller-Ribeiro et al. (2016), suggest that the behavioral, autonomic and respiratory responses generated by the caudolateral superior colliculus in response to external salient stimuli are mediated, at least in part, by direct descending inputs to select brainstem nuclei, and do not depend upon connections with forebrain regions (such as the medial prefrontal cortex, amygdala, and hypothalamus) that are known to play important roles in generating defensive responses to psychologically stressful stimuli (Bondarenko et al., 2015; Carrive, 1993; Dampney, 2015; McDougall et al., 2004; Mohammed et al., 2016; Ulrich-Lai and Herman, 2009).

## Acknowledgments

This work was funded by the National Health & Medical Research Council (APP2001128, APP1127817), the Hillcrest Foundation (IPAP2019_0933, IPAP2018_0437) and by Macquarie University. EL, BD & CS were Macquarie University postgraduate scholars. The authors are grateful to Pascal Carrive, for providing access to his precious thermal camera with patience and grace, and to Nick Everett for the use of his ultrasonic microphone.

## Author Contributions

EL: stereotaxic injections & optrode implantation, behavioral, plethysmography & electrophysiological experiments, data analysis & manuscript draft. BD: anterograde tracing, imaging & analysis, drafting of results. CS: Trans-synaptic tagging of GiA bulbospinal neurons, histology, imaging & validation. EM: behavioral experiments, tail thermography & analysis, drafting of results, histological reconstruction & imaging. AT: stereotaxic injections, optrode implantations & electrophysiological recordings. PB: EEG & plethysmography experiments. AA: AAV-CBA-tdTomato design & construction, drafting of the manuscript. RD: Conceptualization, experimental design, interpretation, drafting of the manuscript and funding. CH: conceptualization & interpretation. JC: supervision and interpretation of behavioral & USV recordings, funding. AKG: conceptualization, supervision & interpretation of tracing experiments, drafting of the manuscript & funding. SM: conceptualization, supervision of electrophysiology experiments, data analysis, drafting of the manuscript & funding. All authors edited the manuscript.

## STAR Methods

*Key Resource Table*

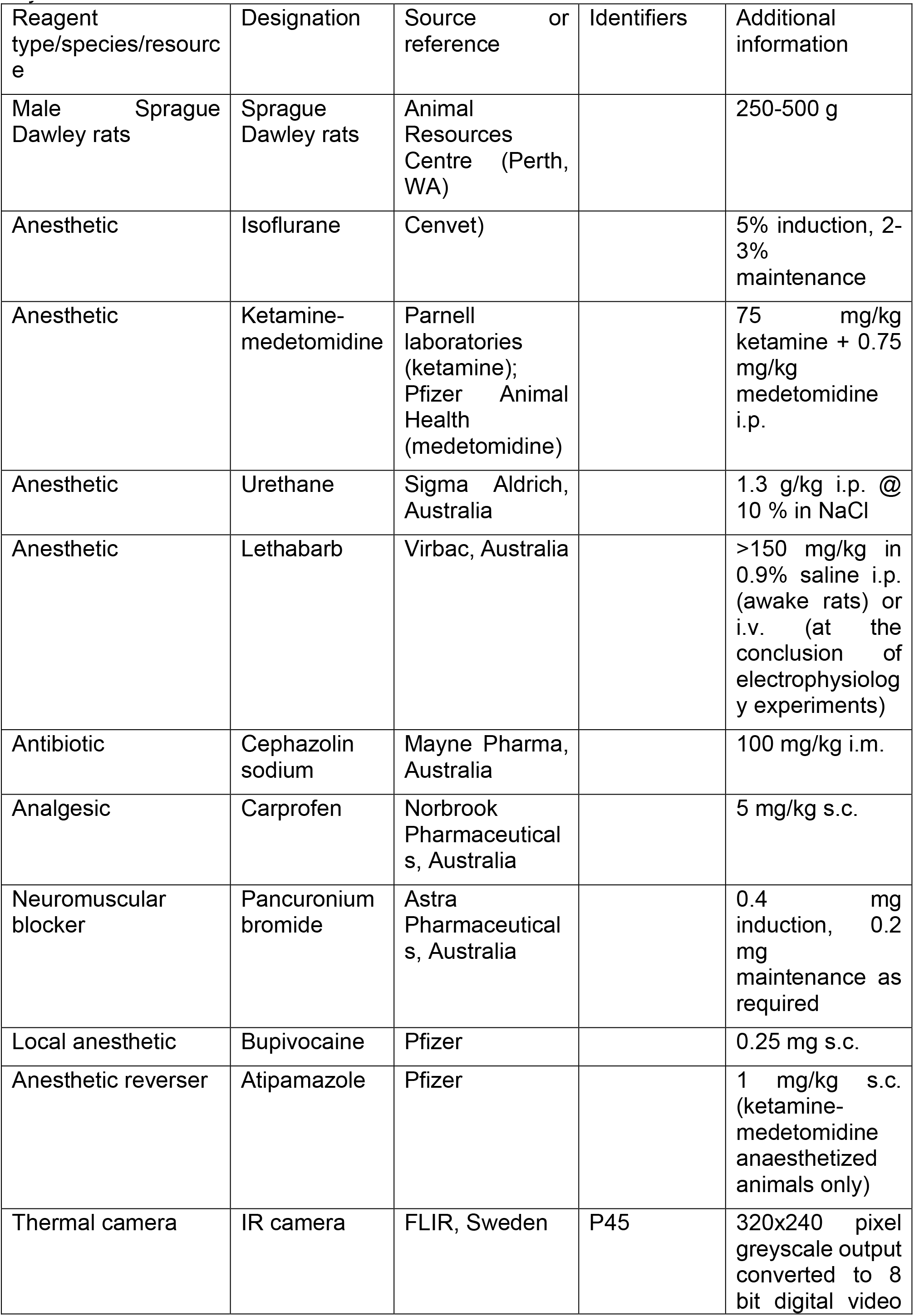
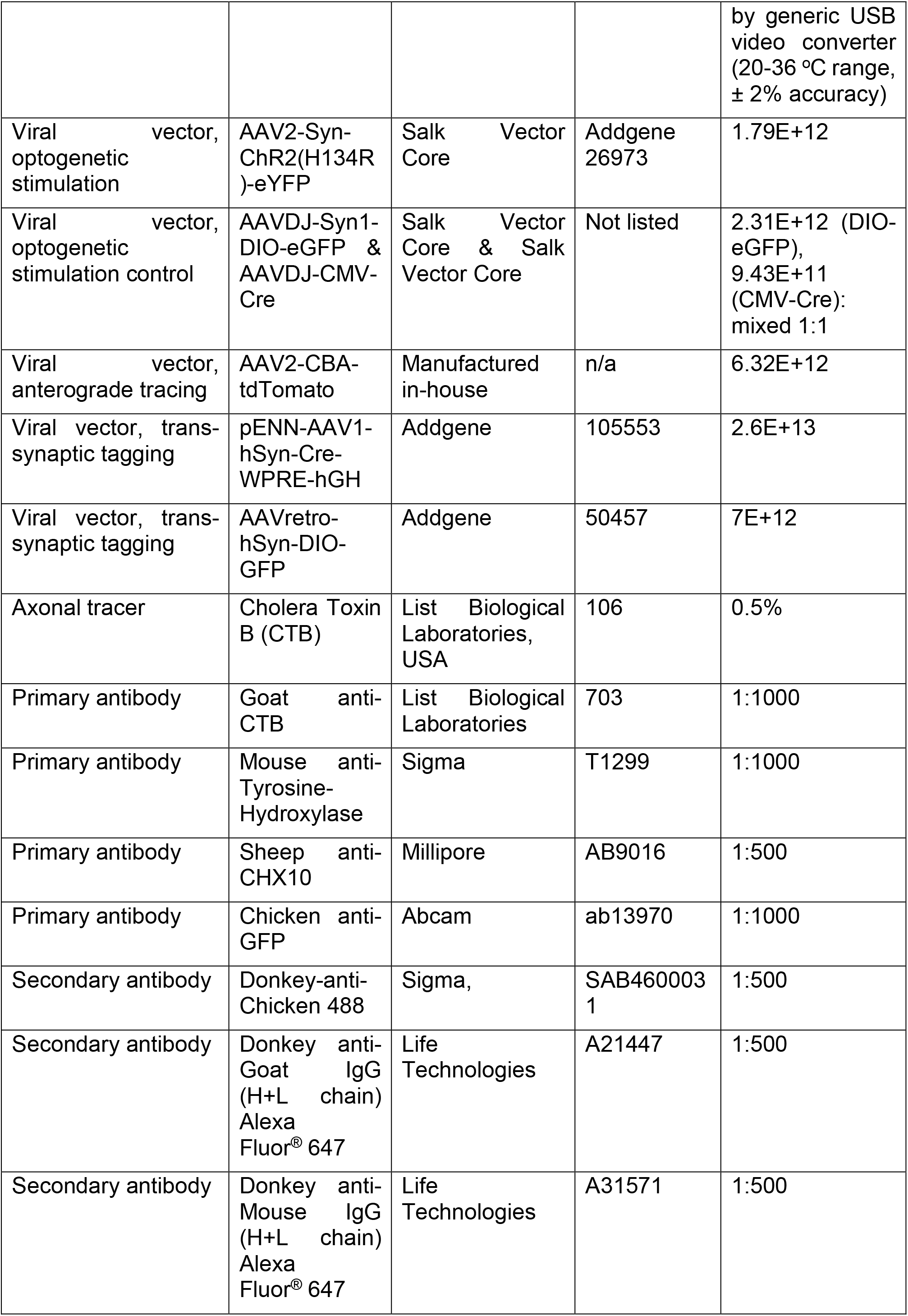

Experiments were approved by the Macquarie University Animal Ethics Committee and conformed to the Australian Code of Practice for the Care and Use of Animals for Scientific Purposes 2013. Adult rats were housed at the Macquarie University Central Animal Facility in groups of two in individually ventilated cages with environmental enrichment materials. Food and water was available *ad libitum*, cages were kept in a temperature and humidity controlled room (21 ° ± 2 °C, 60 % humidity) with a fixed 12h light/dark cycle.

### Animal preparation – recovery experiments

#### dSC vector injection & optrode implantation

Rats were anesthetized with ketamine-medetomidine or isoflurane in oxygen and treated with prophylactic antibiotic, local anaesthetic and analgesic drugs. Anesthetic depth was monitored by assessing respiratory and motor responses to a firm hind paw pinch. Unilateral injections of viral vectors (200-300 nl) were made at the cl-dSC (2 mm lateral, 0.5 - 1 mm rostral, 4.5 – 5 mm deep to Lambda) using borosilicate pipettes under the control of a Neurostar stereotaxic robot. Pneumatic injections were made over 10 minutes and pipettes kept in position for 5 minutes prior to withdrawal.

For cl-dSC stimulation experiments custom-made (Sparta et al., 2012) 0.22 NA 200 μm optical fibers in 2.5 mm ceramic ferrules (‘optrodes’) were inserted 300 μm dorsal to AAV2-Syn-ChR2(H134R)-eYFP or control vector (AAVDJ-Syn1-DIO-eGFP & AAVDJ-CMV-Cre) injection sites and secured to the skull with dental acrylic. Transmission efficiency was noted prior to implantation and used to calibrate laser output (LSR473NL, Lasever Inc) to yield 10 mW maximal output during 20 ms light pulses, equivalent to 320 mW/mm^2^. In a cohort of animals electroencephalographic (EEG) signals were acquired via Teflon-coated silver wires (0.005”, #786000, A-M systems) wrapped around stainless steel jewelers’ screws (1 mm diameter) that anchored the acrylic cap (Burke et al., 2014) and exteriorized via a 6-pin board-board connector header (SPC20500, Element14.com.au).

#### Spinal tracer injection

Three weeks after cl-dSC injection of AAV-CBA-tdTomato a subset of rats received injections of CTB in the T2 spinal cord at coordinates that correspond with the interomediolateral cell column, as described previously (0.75 mm lateral, 1 mm deep: Turner et al., 2013). The spinal processes of ketamine-medetomidine anaesthetized rats were clamped to maintain the spine in a horizontal and elevated position, the T2 spinal cord was exposed, the dura punctured and two 200 nl injections made on each side over a period of 5 minutes before the pipette was slowly retracted. The exposed spinal cord was irrigated with sterile physiological saline, covered with oxidized cellulose hemostat, the wound closed and anesthesia reversed. Post-operative care and monitoring were as described below and rats were perfused 5 days later.

A second group of rats received cl-dSC injections of AAV1-hSyn-Cre (200 nL) under isoflurane anesthesia and bilateral T2 spinal cord injections of AAVretro-DIO-GFP (as above, 100 nl per site). Rats were perfused 3 weeks after surgery.

#### Experimental protocol: behavioral experiments

Rats were habituated to optrode cleaning, handling, and the attachment of fiber optic patch cables for a minimum of three sessions prior to behavioral testing, starting 1-2 weeks after optrode implantation, and also to the rooms used for tail thermography (but not those used for open field test (OFT) or elevated plus maze (EPM) trials). Optrodes were cleaned with 80% ethanol and lubricated with WD-40.

The general effects of unilateral dSC stimulation were investigated in an initial cohort of rats using a low power 470 nm LED system (DC4100 driver with M470F3 light source, ThorLabs, 1-3 mW output). The home cage was moved into a closely fitting high-sided black wooden box, the lid, conspecifics, bedding and environmental enrichment removed, the optrode cleaned and lubricated, and rats left to habituate for 15 minutes. A patch cable fitted with a rotary joint was then attached to the fiber optic ferrule and suspended from an overhead arm and rats were allowed to habituate for a further 15 minutes. Trains of dSC stimulation were completed at 5, 10 or 20 Hz using 20 ms light pulses. Sessions were recorded by video and qualitatively analyzed to categorize the range of responses seen in ChR2 and control rats. Responses were consistent with those reported in response to stimulation of the same region of the dSC in mice, and so were not quantified in detail (Isa et al., 2020).

#### Open Field Test, Elevated Plus Maze & Ultrasonic Vocalization

The OFT, EPM and incidence of spontaneous ultrasonic vocalization (USV) were used to assess whether cl-dSC stimulation was associated with behavioral correlates of anxiety-like behaviors at intensities that caused motor effects.

OFT experiments were conducted during the late part of the rats’ light phase in a room that was novel to rats. The apparatus was a 1 × 1 m white box with 40 cm high walls, illuminated overhead (~400 Lumens). An infrared video camera was mounted overhead to record behavior (Motmen Tracker, Motion Mensura, Sydney). The optrode was cleaned and lubricated and the rat was habituated to the room for 15 minutes in the home cage (open lid, home cage contained in closely fitting high-sided black wooden box) before patch cable attachment. The rat was habituated for a further 15 minutes prior to cl-dSC stimulation (320 mW/mm^2^, 1 Hz, 20 ms pulse width, 2 minutes) after which the patch cable was disconnected, the rat placed in the apparatus, and the operator left the room for the remainder of the 15 minute trial. OFT videos were scored using AnyMaze software (Stoelting). The OFT arena was cleaned with 70% ethanol between trials. The percentage of time spent in a 20 cm square box in the center of the testing arena was compared between ChR2 and control animals by unpaired t-test. Graphpad Prism 7 was used for all statistical analysis.

EPM experiments were conducted during the late part of the rats’ light phase in a laboratory that was novel to rats using an opaque grey Perspex cross-shaped apparatus raised 65 cm above the floor and illuminated overhead (~400 Lumens). The arms measured 50 cm long and 10 cm wide, with the closed arms surrounded by a 40 cm high wall. An infrared camera suspended above the apparatus recorded animal movement. The EPM was cleaned with 70% ethanol between trials.

The optrode was cleaned and lubricated and the rat habituated to the laboratory for 15 minutes in the home cage within a high-walled black wooden box before patch cable attachment and a further 15 minutes habituation before photostimulation (320 mW/mm2, 1 Hz, 20 ms pulses, 1 min) The patch cable was then disconnected and the rat placed on the EPM for five minutes with the operator outside the room. Videos of EPM exploration were scored using AnyMaze software. Parameters measured were time spent in closed and open arms and at the far ends of open arms. The percentage of time spent in each zone was compared between ChR2 and control animals by unpaired t-test. The EPM was cleaned with 70% ethanol between trials. USVs were recorded before and during dSC stimulation and during exploration of the EPM using a M500 USB microphone (10-210 kHz, Pettersson Elektronik, Uppsala, Sweden) and sampled at 500 kHz with 16-bit resolution using BatSound Lite (Pettersson Elektronik). Ultrasound recordings were imported to Spike 2 v7.20 (Cambridge Electronic Design, UK) and converted to spectrograms to enable visualization of USVs, identified using criteria described by Wright et al. (2010) and Brudzynski (2013), and categorized as low- or high-frequency (18-30 and 40-90 kHz respectively). Analysis was completed blind to treatment groups. Numbers of low- and high-frequency USVs were compared between ChR2 and control groups using unpaired t-test.

#### Plethysmography and electroencephalography (EEG)

To assess the effects of dSC stimulation on respiratory frequency and EEG activity rats were habituated to optrode cleaning, patch cable and EEG lead attachment, and a custom-built positive-pressure whole-body plethysmography chamber constructed from a 5.35 liter glass desiccating jar fitted with an overhead airtight port for passage of cabling. Respiratory frequency was extracted from the peaks of DC-removed, bandpass filtered (0.1-20 Hz) recordings of chamber air pressure, captured using Spike 2 using a Power1401 ADC (Cambridge Electronic Design). EEG was amplified and bandpass filtered (0.1 −100 Hz, CME BMA 400) and simultaneously recorded. Stimulation parameters were titrated such that 10 s trains of 5 – 10 Hz dSC stimulation evoked minimal motor effects (e.g. head turning), avoiding gross motor effects that confound the interpretation of plethysmographic recordings. Trials were conducted with rats in quiet wakefulness with at least 5 minutes between stimuli.

Respiratory effects were quantified by comparing average respiratory frequency in the 10 s preceding stimulation to the maximal respiratory frequency recorded during stimulation (running average, 3 s time constant). Significant differences in effect size between control and ChR2 animals were identified using Mann Whitney test. Fast Fourier Transformations of EEG activity recorded 10 seconds prior to and during dSC stimulation were constructed in Spike 2 (Hanning window, 2048 Hz), normalized with respect to total power recorded during baseline, and compared using repeated measures 2-way ANOVA.

#### Tail thermography

Rats were habituated to patch cable attachment and the behavioral laboratory prior to testing. During the light phase of the test day rats were individually transported to the laboratory (ambient temperature 23 °C), nesting removed, and the home box placed within a high-walled black wooden box with an IR camera 1 m overhead. Rats were allowed to habituate in low lighting for 15 minutes before optrode cleaning, lubrication and attachment to a rotary-coupled patch cable, followed by further habituation for at least 15 minutes or until tail temperatures stabilized as indicated by thermography. Video recording and stimulation were initiated using an automated digital sequence to record 10 minutes baseline behavior, 10 minutes 1 Hz dSC stimulation (320 mW/mm^2^, 20 ms pulse width) and 10 minutes post-stimulation activity.

Tail temperature was quantified from IR video images sampled at 10 s intervals. For each image a ROI was manually drawn around the tail and temperature estimated from the greyscale intensity of the brightest (i.e. warmest) pixel (black = 20°C, white = 36°C) using ImageJ Fiji. Each datapoint was expressed relative to the average temperature recorded during the baseline period for that animal and smoothed (5 point rolling average). Significant differences in the effects of photostimulation on tail temperature in ChR2 and control animals were identified using RM 2-way ANOVA. Recordings with unstable tail temperature during the baseline period were excluded from analysis.

The effects of intermittent stimulation of the left dSC on motor behavior was extracted from the same videos: the number of complete rightward circles made during stimulation was counted; significant differences in circling behavior between ChR2 and control animals were detected using Student’s t-test. The distance traveled before, during and after stimulation was measured using Tracker software (Open Source Physics): videos were downsampled to 1 frame/second and the position of the nose manually marked. The relative nose-position from frame-to-frame was smoothed (9 frame rolling average) and the effects of time and dSC stimulation determined by RM 2-way ANOVA.

### Terminal electrophysiology experiments

#### Physiological effects of dSC stimulation

Rats were anesthetized with urethane, placed on a thermostatically controlled heating pad (Harvard apparatus, MA, USA: core temperature 36.5 – 37.5 °C), the flank shaved, and the carotid artery and jugular vein cannulated to enable monitoring of arterial blood pressure (AP) and provide intravenous access respectively. The trachea was intubated with a 14 g cannula, and the rat placed in a stereotaxic frame. Anesthetic depth was monitored throughout surgery via assessing respiratory, pressor and motor responses to firm hind paw or tail pinch. Supplementary urethane (10% initial dose, i.v.) was administered as required.

The postganglionic splanchnic sympathetic nerve was isolated via a retroperitoneal approach, mounted on bipolar silver electrodes and embedded in silicon polymer (Silgel®,Wacker Chemie AG). Central respiratory activity was monitored via direct recording of the phrenic nerve or diaphragmatic electromyogram (EMG), recorded via Teflon-coated steel wire electrodes (Le et al., 2016). Signals were amplified, bandpass filtered (100-1000 Hz, BMA-400, CWE Inc.) and sampled at 3000 - 5000 samples/s (Spike 2). In most cases experiments were conducted in spontaneously breathing animals, supplemented with oxygen; in 4/19 experiments rats were vagotomized, paralyzed with pancuronium bromide and artificially ventilated.

In experiments that measured physiological effects of dSC-GiA terminal activation rats were placed in a nose-down position and the dorsal medulla exposed. The caudal pole of the left facial nucleus was electrophysiologically mapped using a 1 MΩ glass pipette containing 3 M KCl and a 200 μm fiber optic positioned 0.5-0.75 mm lateral to midline, 300 μm ventral to the facial nucleus at rostrocaudal coordinates corresponding to the caudal margin of the facial nucleus.

To assess the effects of 10 or 20 Hz stimulation of the dSC or dSC-GiA terminals, systolic AP and rectified SNA were smoothed (1 s time constant), downsampled to 10 Hz and exported in contiguous blocks that started 20 s prior to and ended 30 s after the onset of stimulation; changes in systolic AP were expressed relative to the average baseline value; SNA was normalized relative to baseline (100%) and noise (0%: obtained by hexamethonium bromide at the conclusion of experiments: 5 mg, Sigma Aldrich, Australia). Significant differences between ChR2 and control animals were identified by RM 2-way ANOVA. Heart- and respiratory rate were derived from systolic AP and the leading edge of diaphragmatic EMG/ phrenic neurograms burst respectively and were quantified by measuring the maximum deviation of the smoothed running averages (bin width 1 and 3 s respectively) from baseline. Statistically significant differences in responsiveness of ChR2 and control animals were detected using unpaired t-test. Ensemble averages of rectified smoothed (τ = 20 ms) SNA responses to dSC or dSC-GiA terminal stimulation were compiled from 200-600 consecutive stimuli delivered at 0.5 Hz.

#### GiA neuronal responsiveness to dSC stimulation

Rats were prepared as described above, bilaterally vagotomized, and artificially ventilated in oxygen-enriched room air under neuromuscular blockade. The T2 spinal cord was exposed and a bipolar concentric stimulating electrode (Rhodes NE-100) aimed at the dorsolateral funiculus (to antidromically activate the axons of bulbospinal neurons) at coordinates that resulted in maximal SNA responses to 200 μA cathodal pulses (0.2 ms). The facial nucleus was mapped as described above and extracellular recordings of spontaneously active neurons made via glass microelectrodes filled with 0.9% NaCl (30-50 MΩ), amplified and filtered (Axoclamp 900A, Molecular Devices, Palo Alto, CA, 300–3,000 Hz) and sampled at 10,000 samples/s using Spike 2. The region corresponding to the GiA (0.4-0.9 mm from midline, 2.8 – 4.1 mm deep to the dorsal surface, ±0.4 mm rostrocaudal to the caudal pole of the facial nucleus) was surveyed. Spontaneously active neurons with spinal projections were identified by constant latency antidromic responses to spinal stimulation that collided with spontaneous orthodromic spikes (Lipski, 1981). Putative sympathetic premotor neurons were functionally identified from systole-triggered averages of neuronal discharge or spike-triggered averages of SNA, defining features of other sympathetic premotor populations (Chen and Toney, 2010; Kanbar et al., 2011; McMullan et al., 2007). Spike-triggered averages of rectified & smoothed (50 ms time constant) SNA were generated from 400-26,000 spike-triggered sweeps. ‘Dummy’ spikes that occurred at the same average frequency as a negative control (Morrison et al., 1988; Turner et al., 2013).

Neuronal discharge was plotted as mean frequency (2 s time constant) and effects of optical cl-dSC stimulation at 10 or 20 Hz expressed as maximum change in discharge during stimulation compared to the average discharge in the 10 preceding seconds. Responses to low-frequency cl-dSC stimulation were identified from peristimulus time histograms constructed from 200-400 consecutive trials.

#### Euthanasia, perfusion, histology & imaging

At the conclusion of experiments animals were overdosed with sodium pentobarbital and perfused with 500 ml heparinized saline followed by 500 ml 4% paraformaldehyde. Brains were removed, post-fixed at room temperature overnight, and cut in 50 μm thick coronal sections using a vibrating microtome (Vt1200S, Leica) at 200 μm intervals.

To verify dSC injection sites sections were mounted immediately onto glass slides with mounting medium (Dako Fluroshield with DAPI, Sigma), cover-slipped, and native reporter fluorescence imaged using a Zeiss Z2 epifluorescence microscope at 10x. Pilot experiments indicated weak or absent behavioral responses from off-target injections: 12 successful experiments were chosen to generate a heat map demarcating reporter expression in experiments associated with typical behavioral responses. To construct the heat map three greyscale images were used from each animal; one that contained the center of the injection site, one 200 μm rostral to the center and one 200 μm caudal to the center. Images were downsampled to 25%, Gaussian blurred (6 pixel) and eYFP fluorescence normalized so that the value of the brightest labelling became white and the background became black. Images were aligned to the appropriate rat atlas plates (Paxinos and Watson, 2006) and individual images from each bregma level overlaid and averaged using ImageJ. Normalized averaged greyscale images were then colorized using the ImageJ Thermal LUT.

‘On target’ experiments were defined as having reporter expression distributed 6.8 to 7.8 mm caudal to Bregma, mostly within the deep grey and deep white layers of the superior colliculus, medial to the nucleus of the brachium of the inferior colliculus, dorsal to the precuniform area, and lateral to the dorsolateral and lateral periaqueductal grey. If YFP reporter expression or optrode tracks fell outside these boundaries data were excluded from analysis.

#### Immunohistochemistry

Brain sections were permeabilized in for 3 × 15 minutes in TPBS with 0.1 % Tween20 or 0.2% Triton-100 and blocked for nonspecific binding in TPBS containing 2% bovine serum albumin and 0.2% Triton-100 for 1 h at room temperature. Primary antibodies were added to blocking buffer and incubated for 48 hrs at 4°C. Sections were then washed in PBS 3 × 30 minutes and incubated in TPBSM with 5% NHS and secondary antibodies for 12 hrs at 4°C. Sections were washed again for 3 × 20 minutes in PBS before being mounted on glass slides with Dako Fluroshield mounting medium with DAPI and cover slipped.

Fluorescence imaging was performed on a Zeiss Z2 epifluorescence microscope or Leica TCS SP5X confocal microscope. Images were prepared and analyzed using ImageJ Fiji and Corel Photopaint X7.

